# Interplay Between Pulmonary Membrane Properties and Lung Disease: A Study of Seven Bottlenose Dolphins

**DOI:** 10.1101/2025.10.17.683140

**Authors:** Marilyn Porras-Gómez, Bengu Sueda Sengul, Nurila Kambar, Sari Gluck, Kristen Flatt, Celeste Parry, Carolina Ruiz Le-Bert, Diego Hernández-Saavedra, Cecília Leal

## Abstract

Pneumonia is the leading cause of morbidity and mortality of bottlenose dolphins *Tursiops truncatus*. We investigate a series of rare and opportunistic samples of pulmonary surfactant membranes (PSMs) extracted from lungs of seven dolphins in the care of the U.S. Navy Marine Mammal Program. We found a striking correlation between PSM structure, lipidome, and mechanical properties with the severity of lung injury. Specifically, lipidomics reveals exacerbated contents of cardiolipins, confirming a result obtained for terrestrial mammals afflicted by experimental pneumonia. Employing a battery of X-ray scattering, atomic force, and electron microscopy, we evaluate how the altered lipid composition impairs the structural integrity of the PSM and leads to dehydration and enhanced rigidity. Our findings demonstrate that the function of pulmonary surfactant membranes goes far beyond lowering alveolar surface tension, regulating their biochemical and biophysical properties with lung pathology progression. This knowledge will be useful to the development of future diagnostics and therapeutics of respiratory diseases targeting lung membranes.

## Introduction

The mammalian lung evolved to enable highly efficient gas exchange in the alveoli while mechanically undergoing repeated expansion and contraction during breathing. This process is particularly efficient in diving mammals that have cartilaginous airways extending to the mouth of the alveolar sacs and conforming lungs that allow: i) movement of air during lung compression, ii) shutting down of N_2_ adsorption and gas exchange, and iii) alveolar collapse. ***Denison et al. (1971)*** While humans exchange about 20% of their lung capacity with each breath, dolphins take short and deep breaths, with an exchange of 75% to 90% of air in one-third of a second. ***Fahlman et al. (2015***, ***2017***) These breathing adaptations put them at increased risk of respiratory infections by enabling deep lung exposure to airborne threats at the marine surface. Pneumonia is an inflammatory respiratory disease that primarily affects alveoli and is the primary cause of morbidity and mortality in marine mammals. ***Venn-Watson et al. (2012)***; ***Díaz-Delgado et al. (2018)***

Central to facilitating gas exchange, ***Olmeda et al. (2010)***; ***Kim et al. (2020)*** lung innate immunity, ***Chroneos et al. (2010)*** and lung mechanics ***Prokop and Neumann (1996)***; ***Hermans et al. (2015)*** is the alveolar air/aqueous hypophase lining. This consists of a phospholipid-rich monolayer that reduces the surface tension in the alveoli to 5–20 mN/m, and that is why it’s called *lung surfactant*. In reality, this interfacial monolayer directly connects to a complex and dynamic lipoprotein system occupying the whole volume of the alveolar hypophase. An updated squeeze-out model describes the monolayer interconnected to tubular membranous structures (tubular myelin, TM) ***Xu et al. (2020)***; ***Keating et al. (2012)***; ***Zuo et al. (2008)***; ***Zhang et al. (2011b)*** stemming from lipoprotein multilamellar bodies (MLBs) that are secreted by Type II pneumocytes. ***Lettau et al. (2022)***; ***Ochs (2009)***; ***Olmeda et al. (2017)***; ***de la Serna et al. (2004)***

Lung surfactant irregularities have been associated with a variety of lung pathologies, and it should be expected that its composition, structure, and mechanical properties will change in response to lung injury. ***Milad and Morissette (2021)*** Indeed, neonatal respiratory distress syndrome (NRDS) is characterized by insufficient pulmonary membrane production, whereas pulmonary alveolar proteinosis leads to pulmonary surfactant accumulation in the alveoli ***Milad and Morissette (2021)***; ***Dushianthan et al. (2011)***; ***Wang et al. (2021)*** leading to impaired surface activity and difficulty breathing. Mutations in pulmonary membrane-related genes (i.e., biogenesis and regulation) have been identified as an important cause of severe pulmonary diseases ***Whitsett et al. (2015)***; ***van Moorsel et al. (2021)*** and even impact nonspecific lung pathologies such as lysosomal storage diseases. ***Paget et al. (2021)*** Moreover, pulmonary membrane composition also plays a role in the diagnosis of lung diseases. Levels of specific surfactant proteins, for example, may have utility as prognostic and diagnostic indicators in patients with interstitial lung disease. ***Kaieda et al. (2020)***; ***Takahashi et al. (2006***)Recently, two lung surfactant proteins (cetacean surfactant protein D and lipopolysaccharide-binding protein) were identified as potential biomarkers of pneumonia in bottlenose dolphins. ***Hamano et al. (2023)***

The pulmonary surfactant membrane lipidome, however, is less explored for hallmarks of lung disease and diagnosis, despite evidence that pneumonia affects levels of glycerolipids, glycerophospholipids, and phosphatidylglycerol. ***Shan et al. (2018)*** Nancy Ray and co-authors ***Ray et al. (2010)*** reported that pneumonia leads to a change in pulmonary membrane phospholipid composition in several terrestrial mammal species, specifically to an accumulation of a mitochondrial phospholipid, cardiolipin, in lung fluid. We have previously shown that excess cardiolipin compromises the structure and elasticity of pulmonary membranes, leading to gas exchange imbalance. ***Kim et al. (2020)***; ***Porras-Gómez et al. (2022b)*** Disruption of membrane and tissue elasticity affects alveoli compressibility ***Petty et al. (1979)*** and increases alveolar surface tension, leading to atelectasis ***Günther et al. (2001)*** and ultimately ineffective breathing. ***Günther et al. (2001)***; ***Petty et al. (1979)*** In this paper, we examine how the severity of respiratory diseases correlates with the lipidic composition, structure, and mechanics of pulmonary surfactant membranes extracted from Navy bottlenose dolphins (*Tursiops truncatus*) exhibiting a range of respiratory pathologies, including pneumonia. These rare, retrospective, and opportunistic samples come from a large biobank dating back decades and belonging to the U.S. Navy Marine Mammal Program. Lung exudates include pulmonary extracts and lung tissue. We found that cardiolipins are predominantly more abundant in abnormal dolphin lung exudates, leading to a loss of elasticity in membranes and lung tissue. We also noted that plasmalogen phosphocholines, a phospholipid class that protects against oxidative damage, appear more abundant in lung membranes with a recent history of pneumonia compared to chronically diseased lungs and may elucidate an association of plasmalogen activity with early disease stages. We evaluate membrane fluidity and organization and present a link between lateral membrane heterogeneity and the physiological function of native pulmonary membranes under pathological conditions. In summary, we establish strong associations between the lung disease and the biophysical properties of pulmonary membranes extracted from bottlenose dolphins. This knowledge is central to the development of new prevention, diagnosis, and treatment strategies of pulmonary diseases, as well as the potential for repurposing and enhancement of pulmonary membrane replacement therapies. To the best of our knowledge, this is the first study that links pulmonary membrane homeostasis with pulmonary disease in marine mammals.

## Results

### Lung extracts reproduce the length-scale and structural diversity of native pulmonary membranes

Bronchoalveolar lavages (BALs) were performed as a minimally invasive routine procedure to diagnose Navy dolphins affected by a range of naturally occurring respiratory pathologies, i.e., illness and pathology were not artificially induced. We performed organic extractions to separate the lipid material from BAL lung exudates. Fig. 1A depicts the reconstitution of BAL extraction from a healthy animal. To validate that the extraction procedure enabled the study of membrane suspensions, we performed an organic extraction, which led to the collection of crude amphiphilic molecules (lipids and surfactant proteins). Subsequently, we subjected the samples to ultra-centrifugation cycles that separate debris and remaining cells, including neutrophils, leukocytes, and other immune cells that could have infiltrated the hypophase, especially because pneumonia is characterized by increased neutrophil infiltration. We then rehydrated (schematics of Fig. 1A) the organic phase tagged with a fluorescence lipid probe and inspected the sample by laser confocal fluorescence microscopy (Fig. 1B). We observe that lung lipid extracts reassemble upon rehydration into small, large, single, and multilamellar vesicles. These aggregate forms reproduce the expected membrane microstructures observed in native alveolar hypophase. Higher resolution transmission electron microscopy (TEM) negative staining images of rehydrated BAL lipids show the presence of elongated and multivesicular (MVS) structures (Fig. 1C) consistent with previous studies of native BALs. ***Lettau et al. (2022)***; ***Ochs (2009)***; ***Olmeda et al. (2017)*** This indicates that pulmonary membranes can be successfully extracted and reconstituted from BALs of Navy dolphins. No evidence of cells or cell debris could be detected by confocal, optical, or electron microscopy.

**Figure 1.**
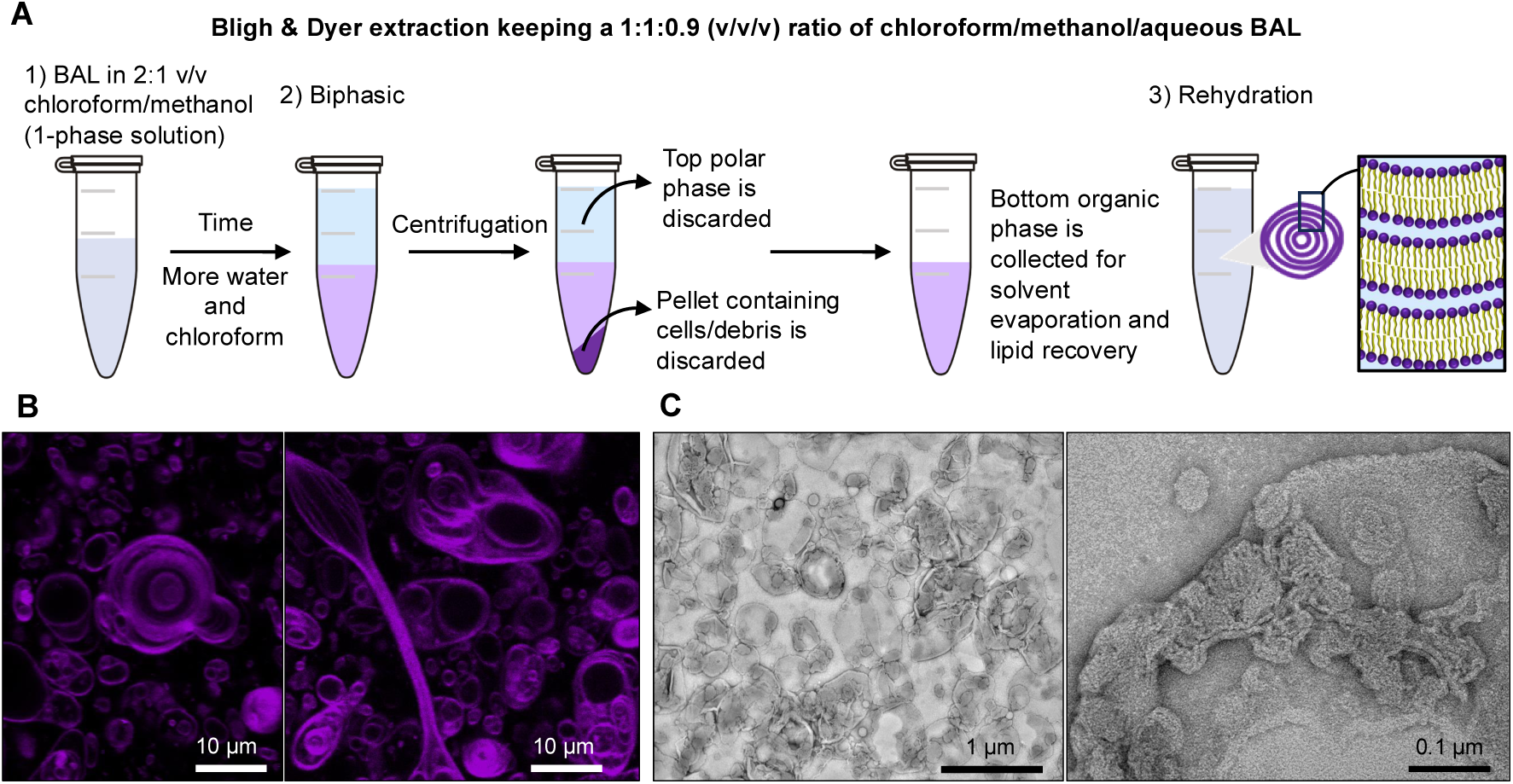
Lung exudate extraction procedure and visualization. A) Schematics of organic extraction to obtain the lipidic phase in the form of a dry film after solvent evaporation and subsequent lipid self-assembly upon rehydration of the dry film in a capillary tube. B) Laser confocal fluorescence microscopy images of fluorescently labeled pulmonary lipid extracts in water. C) Transmission electron microscopy images of BAL lung extracts from a dolphin with clinically healthy lungs exhibiting multivesicular structures (MVS) and tubular structures.

### Lipidome signatures characterize lung injury

Previous studies have compared bronchoalveolar lavage (BAL) lipids between terrestrial mammals (including humans) to those of marine mammals ***Gutierrez et al. (2015)***; ***Spragg et al. (2004)*** such as diving pinnipeds (seals). In general, phospholipid concentrations are greater in BAL from pinnipeds than in humans, with a greater percentage of phosphatidylcholine (PC) compared to other phospholipid species. Pinniped surfactant was significantly enriched in the fluidic PC species. This indicates that diving mammals may have larger surfactant pool sizes and larger aggregates with anti-adhesive properties than surfactant from terrestrial mammals. Comparison of odontocete—dolphins and porpoises—lipids to that of pinnipeds found that odontocetes have comparable PC composition to that of pinnipeds. ***Gutierrez et al. (2015)*** The difference in fluidic PC levels in marine mammal lung surfactant constituents is likely to facilitate film respreading during rapid alveoli collapse. ***Spragg et al. (2004)***; ***Denison et al. (1971)***; ***Miller et al. (2006)***

We categorized lung exudates from dolphins with acute/chronic lung disease or no lung disease at the time of sample collection (Table 1) per diagnosed by the team of U.S. Navy Marine Mammal Program veterinarians. To study the lipidic composition of lung exudates, we performed structural lipidomics and identified ≈490 lipid species (Fig. 2). Lipidomic analysis shows that global lipid profiles help distinguish samples by presence or absence of lung disease, as depicted through the principal linear discriminant analysis (Fig. 2A). The multivariate analysis demonstrates that over 70% of lipid variations can be explained by two components that support disease clustering. More-over, among the top discriminant lipid species for lung severity are several cardiolipins (CL), phosphatidylserines (PS), phosphatidylcholines (PC), and plasmalogens (Fig. 2B). To identify the potential involvement of lipids in biophysical and biochemical processes, we performed a lipid ontology (LION) analysis. Compared to lung extracts with no lung disease, we identified a significant enrichment within components of membranes in the endoplasmic reticulum (ER) and mitochondria, as well as plasmalogens and glycerophospholipids in lungs with acute/chronic disease (Fig. S1). Here we show marine mammals also accumulate excess cardiolipins during pneumonia (Fig. 2C). Alterations in the degree of saturation are also observed with disease, yielding more acyl chains at lower unsaturation levels. Lipidomic profiling confirmed that lung fluid from dolphins with chronic/acute lung disease (Fig. 2D) contains higher levels of a wide variety of cardiolipin species, compared to lungs from dolphins with no lung disease. Lung disease significantly alters lipid signatures in lung exudates.

**Figure 2.**
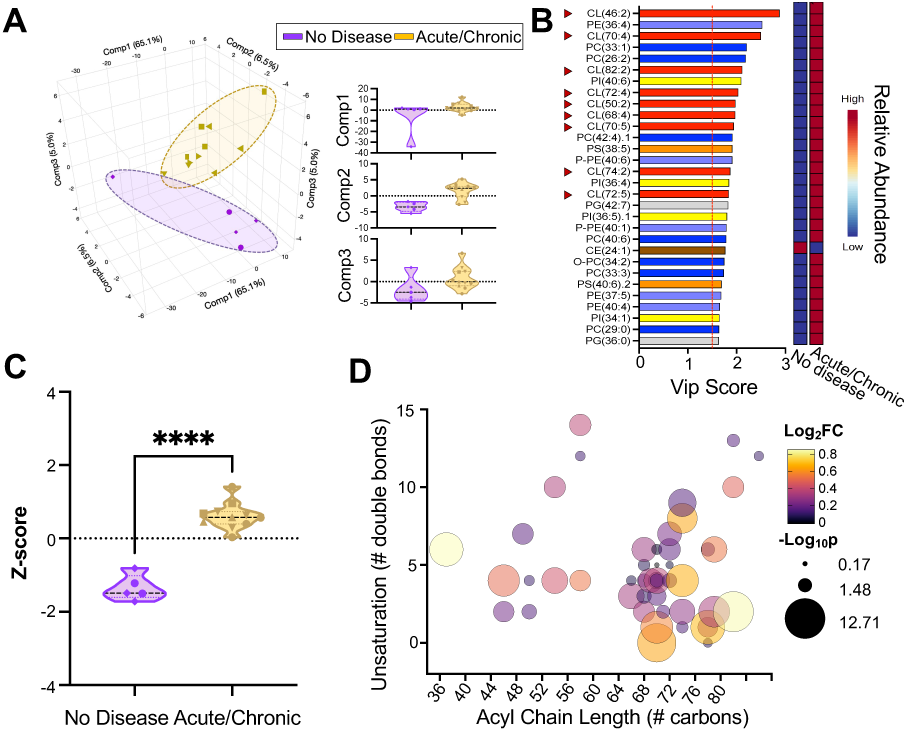
Lipidomic analysis. A) Principal linear discriminant analysis of lipids from lung exudates. Replicates are represented by similar symbols. The values for components 1, 2, and 3 are shown. B) Top 30 species that discriminate between groups and account for variation within component 1. Cardiolipin species are indicated by red triangles. C) Cardiolipin abundances depicted as a Z-score. D) Cardiolipin species analysis in samples from acute/chronic pulmonary infection. Cardiolipin fatty acyl-chain length is shown on the x-axis, and cardiolipin fatty acyl unsaturation level (double bonds) is shown on the y-axis.

**Table 1.** Categories assigned to Tursiops truncatus lung exudates with corresponding number of animal samples (*n*)

| Pulmonary condition category | Description |
| --- | --- |
| *Chronic or acute lung disease, n=5* | History of chronic or acute pulmonary infection such as bacterial or fungal pneumonia, mild alveolar interstitial syndrome, or recent history of pneumonia and recovered with treatment |
| *No lung disease, n=2* | No history of pulmonary disease at the time of sample collection |

| Pulmonary condition category | Description |
| --- | --- |
| Chronic or acute lung disease (n=5) | History of chronic or acute pulmonary infection such as bacterial or fungal pneumonia, mild alveolar interstitial syndrome, or recent history of pneumonia and recovery after treatment |
| No lung disease (n=2) | No history of pulmonary disease at the time of sample collection |

### Lung pathology disrupts lateral membrane domain organization

Pulmonary membranes comprise a fine balance of saturated and unsaturated lipids that laterally phase-separate into membrane domains ***de la Serna et al. (2004***, 2013); ***Dhar et al. (2012)***; ***Alonso et al. (2004)*** that maintain mechanical robustness and rapid re-spreading to reduce surface tension during breathing. ***Postle et al. (2001)*** The observed alterations in lipid composition and saturation (Fig. 2) are likely to impact the lateral organization of lung membrane domains. We analyzed extracted multilayered membranes by Atomic Force Microscopy (AFM) to evaluate morphological changes indicative of phase separation. It is important to emphasize that the samples studied are not adsorbed monolayers, as is most common for AFM studies of lung surfactants, but rather multilayered bilayer stacks (≈100 nm in thickness) of lung membrane extracts that were supported onto solid substrates by spin-coating at near 100% RH. AFM analysis conveys information about the morphology and spatial distribution of phase-separated domains such as lipid *gel*-phase (L_β_) and the *liquid*-phase (L_α_) domains (Fig. S2). The L_β_ phase comprises thicker bilayers with well-packed saturated lipids forming stiffer domains vs. (L_α_) domains, which are thinner and more *fluid*-like. AFM fast force mapping is a valuable complement to topography analysis as it quantitatively evaluates the interaction forces between the probe and the sample. The combination of topography and force mapping AFM demonstrates that locally L_β_ domains are indeed stiffer than the surrounding L_α_ phase and lower adhesion forces. In a similar fashion, local domain mechanical properties have been mapped for DPPC:DOPC supported bilayers with AFM. ***Saavedra V et al. (2020)*** Our data suggests that lung pathology significantly disrupts the morphology of phase-separated L_β_ domains, including size and shape. Quantitative AFM analysis shows that lung disease leads to a reorganization of the immiscible L_β_ -L_α_ membrane domains. The optimization of L_β_ domain area/perimeter (A/P) ratio is indicative of the ability to modulate line tension between the L_β_ and the surrounding L_α_ phases. ***Lin et al. (2007)***; ***de la Serna et al. (2004)*** Lung exudates from dolphins with lung disease conditions display larger and more elongated L_β_ domains (Fig. 3A, right). In contrast, in membranes coming from healthy lungs, the L_β_ phase coalesces into more circular domains (i.e., a wider population around circularity index of 0.8) covering larger surface areas (Fig. 3B-C), indicating the ability to sustain higher line tension.

**Figure 3.**
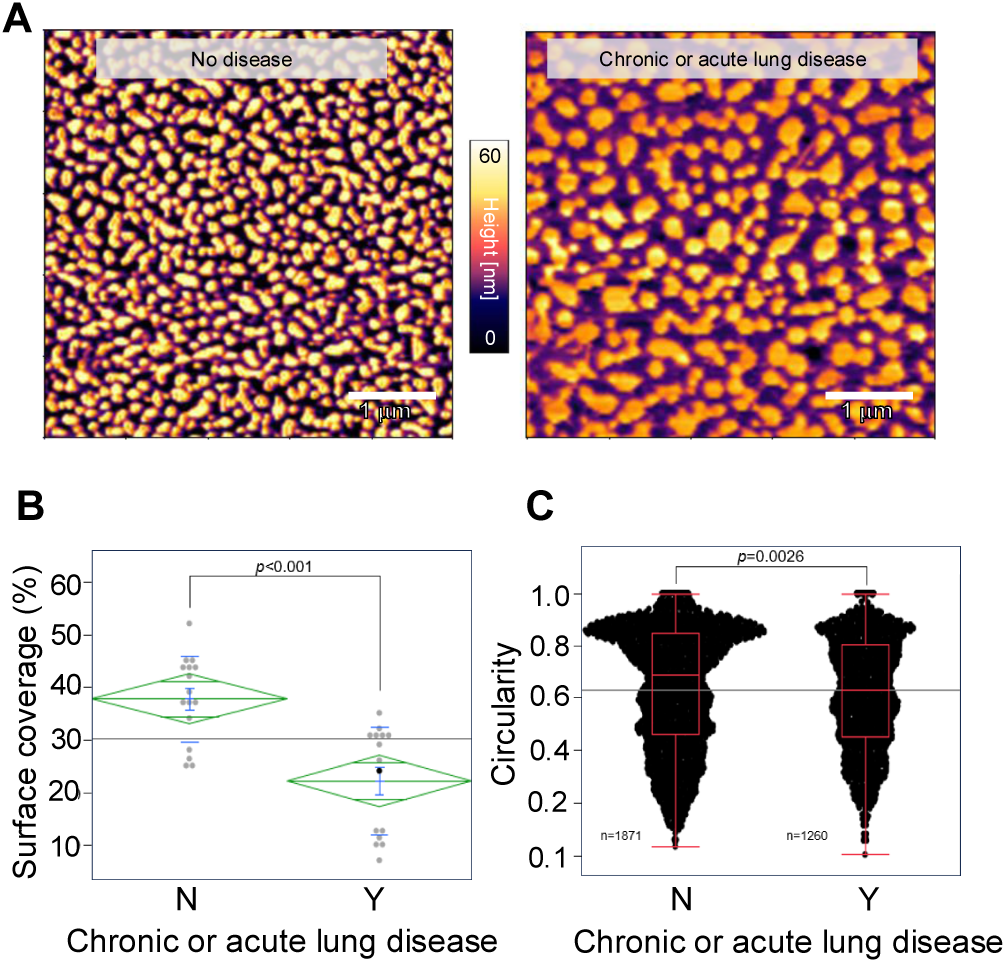
Microscopy analysis of lung extracts and lung tissue. A) AFM topography maps of multilayered lung fluid extracts from dolphins with no disease and with lung disease. B) One-way ANOVA analysis of surface coverage with a significant p-value. C) Oneway ANOVA analysis of domain circularity by lung disease condition, resulting from topography map analysis with a significant p-value. The number of domains analyzed is shown.

### Lung disease modulates membrane and tissue elasticity

We previously employed AFM to demonstrate that pneumonia-associated phospholipids (i.e., cardiolipins) significantly increase the Young modulus of stacked bilayers in both ternary lipid membrane models and bovine lung extracts. ***Porras-Gómez et al. (2022b)*** This stiffening arises because cardiolipin promotes fusion of apposed bilayers within the stacks, which reduces interlayer water content and concomitant higher inter-bilayer adhesion. Consistent with this mechanism, our AFM measurements showed higher adhesion forces in samples with acute or chronic lung disease (Fig. 4A, right), and these effects correlate with elevated cardiolipin abundance detected in samples from acute and chronic lung disease (Fig. 4B). Moreover, a strong positive correlation was observed between cardiolipin abundance and the Young modulus (*R*^2^ = 0.7732; Fig. 4C), indicating induced stiffening of the pulmonary membranes in chronic or acute pulmonary conditions. Notably, stiffening is also observed in lung tissue affected by lung disease (Fig. S3A). Two lung tissue samples from dolphins with lung disease and one from a healthy dolphin postmortem were analyzed with fast force mapping (Fig. S3B-D). In the same way, the Young modulus was higher in diseased tissue. In addition to AFM force mapping, we utilized transmission electron microscopy (TEM) negative staining imaging to investigate the nanoscale morphology of pulmonary membranes in lung tissue (Fig. S3E-G). Lung tissue from dolphins with no lung disease present well-hydrated multilamellar bodies as well as vesicles and well-defined elongated membranes. In contrast, chronic/acute lung disease samples also display well-hydrated multilamellar bodies (Fig.S3F, right) but denser objects and less well-defined membranes can be detected (e.g. all Fig. S3G panel images). Moreover, several membrane forms and dense-core particles of < 100 nm in diameter appear in the lung tissue sample coming from dolphins with lung disease, which could correspond to packed inclusion bodies from alveolar type II cells. ***Have-Opbroek et al. (1990)***

**Figure 4.**
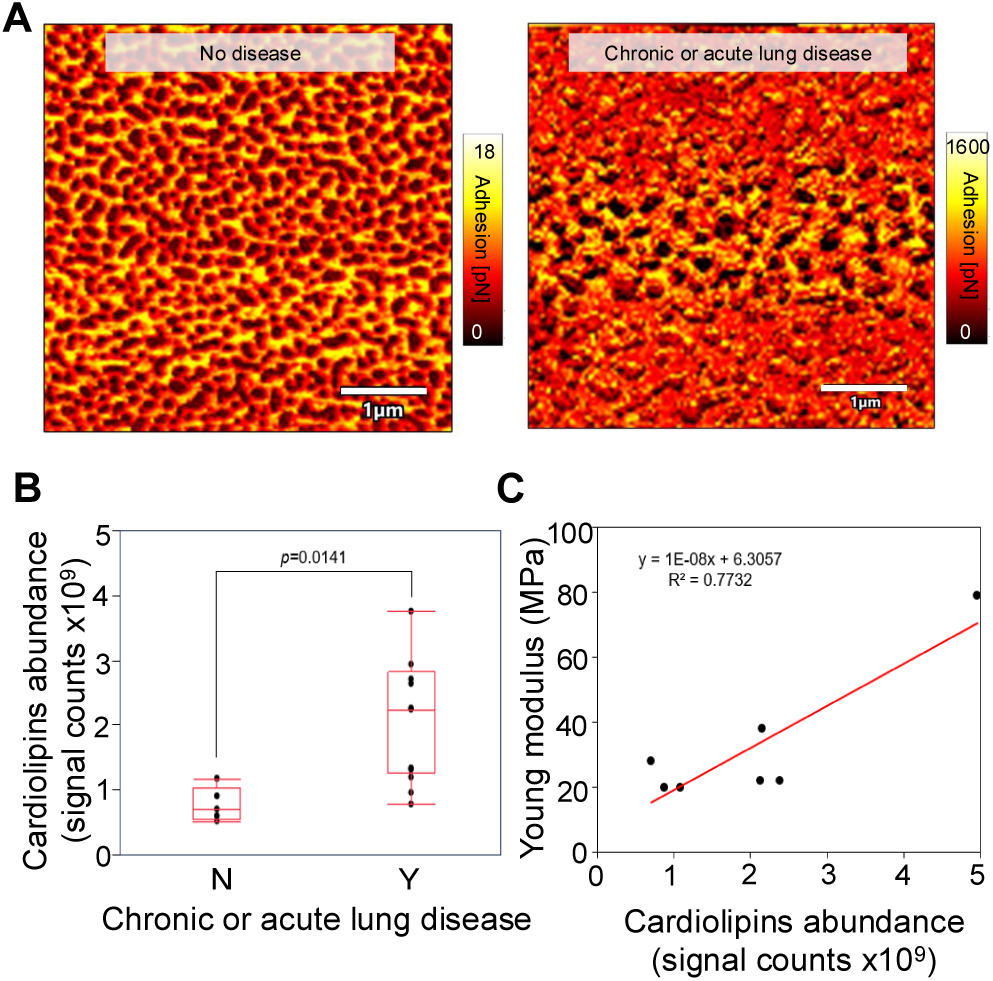
Microscopy analysis of lung extracts and lung tissue. A) AFM adhesion maps of multilayered lung fluid extracts from dolphins with no disease and with lung disease. B) One-way ANOVA analysis of cardiolipin abundance by lung disease condition. Cardiolipins’ abundance values correspond to the normalized signal counts to internal standards and sample weight. C) Bivariate fit of Young modulus by cardiolipin abundance. Linear fit and corresponding equation are shown. Elasticity means of lung extracts were calculated from fast force mapping.

### Lung pathology alters pulmonary membrane ultrastructure

To further understand the effect of disease severity and cardiolipin excess on the structural integrity of pulmonary membranes, we inspected membrane suspensions of lung exudates using small and wide-angle X-ray scattering -SAXS/WAXS (Fig. 5A-C). Fig. 5A-B presents SAXS/WAXS data as integrated intensity (arbitrary units) as a function of wave vector *q* (nm^−1^) for lung extracts from dolphins with and without lung pathologies. Several peaks can be observed in the SAXS regime (Fig. 5A). As shown in Fig. 1, extracted pulmonary membranes can be reconstituted in water, selfassembling into several vesicular and multilamellar bodies (MLBs). SAXS is very sensitive to membrane periodicity at the nanoscale, hence allows for accurate probing of inter-bilayer distances (*d*) in MLBs. The observed Bragg peaks are consistent with a multilamellar phase of pulmonary lipids yielding peak *q*-position ratios of 1:2:3:4(…). This indicates that lung extracts indeed self-assemble into MLBs where bilayers are coherently correlated and separated by a water layer of conserved thickness. In addition to the 100 and 200 Bragg peaks, a scattering form factor envelope is also observed, indicating the presence of vesicular structures. Compared to healthy dolphins, the SAXS pattern is clearly altered for pathological samples. More peaks (up to 500) can be resolved that shift to higher *q* values, and the form factor envelope is less discernible. Similar results were obtained when adding cardiolipin to lung membrane model systems. ***Steer et al. (2018)*** In addition, our lab and others previously found that bovine-extracted pulmonary membranes exhibit a coexistence of two lamellar phases, ***de la Serna et al. (2013)***; ***Larsson et al. (2002)***; ***Steer et al. (2018)*** one rich in unsaturated lipids and one in saturated lipids (0.5-2 nm thicker) that merge at temperatures above 40^◦^C. ***de la Serna et al. (2013)*** The SAXS data of lung exudates present a single lamellar phase even though lipid domains are clearly present, as shown in Fig. 3A and will be later demonstrated by cryogenic electron microscopy (cryo-EM). This is likely because the phase-separated domains are small and not fully aligned in the MLBs.

**Figure 5.**
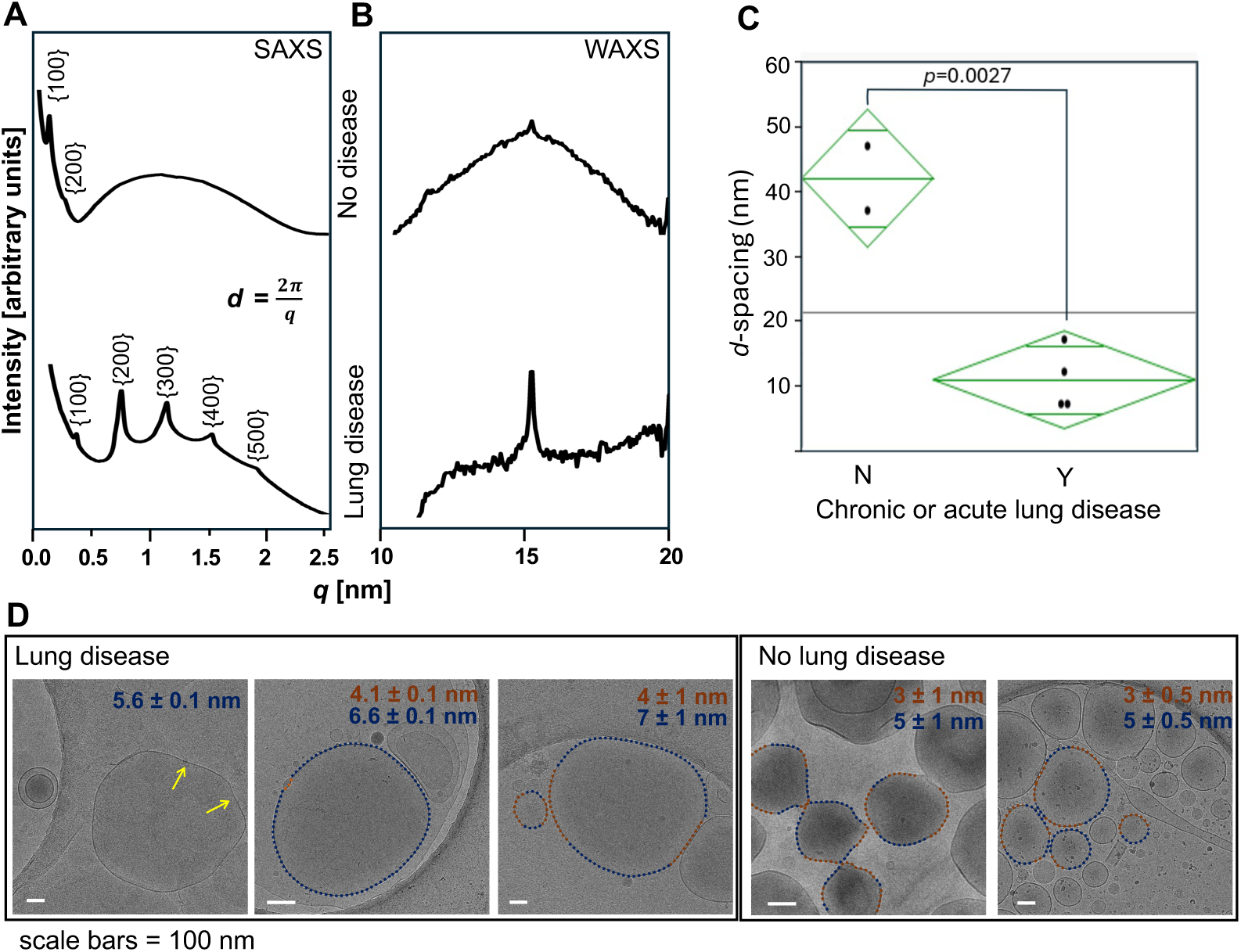
Structural characterization of pulmonary membrane complexes. SAXS (A) and WAXS (B) Intensity vs q data. C) One-way ANOVA analysis of lattice parameter d response to lung disease condition. *d*-spacing was calculated from SAXS peak positions in A. D) Cryo-EM data where scale bars correspond to 100 nm. The Cryo-EM images depict lung extracts from dolphins with chronic or acute lung disease (left) and dolphins with no lung disease (right). Nanodomain size is highlighted in blue (> 5 nm) and in orange (< 5 nm), and was estimated with rim analysis.

The average separation between bilayers in a multilayer stack (*d*-spacing) results from the sum of the lipid bilayer thickness *d*_*B*_ and the water layer thickness *d*_*w*_ and can be calculated from the nth diffraction peak as *d* = *n*2π∕*q*_*n*_. In our previous work, ***Steer et al. (2018)***, when cardiolipin was artificially introduced, we observed that phase separation was exacerbated and that the two multilamellar systems had very different *d*-spacings (Δ*d*> 5 nm), which was attributed to different water layer thicknesses. Fig. 5C plots *d* obtained from membranes extracts calculated from *q*_100_ in Fig. 5A. The bilayer thickness *d*_*B*_ should not vary significantly with lipid composition and was previously estimated to range 5-8 nm ***Porras-Gómez et al. (2022b)*** (see also later Fig. 5D). Hence, drastic changes in *d* likely arise from alterations in the hydration layer *d*_*w*_. SAXS data indicate that lung fluid from dolphins with pulmonary disease exhibits thin water layers (*d* ≈7 nm indicates *d*_*w*_ of 2-6 nm). Conversely, samples from dolphins with no lung diseases exhibit very well-hydrated pulmonary membranes with water layers of *d*_*w*_≈40 nm. These results reinforce that pneumonia-associated membrane alteration leads to aggregation of lipids and dehydration of pulmonary multilamellar bodies. As expected, pulmonary extracts from dolphins with no lung disease exhibit large *d*-spacing, indicating that lung membranes in the alveolar hypophase are fully hydrated, which is imperative for membrane protein function and general lung membrane homeostasis.

WAXS probes membrane organization at the molecular level, i.e., the configurations of lipid fatty acid chains, related to membrane fluidity and acyl chain order. The shape and width of the characteristic WAXS peak at *q* ≈ 15 nm^−1^ for lipid bilayers enable the identification of fluid/gel phases (L_α_/L_β_) (Fig. 5B). For instance, lung extracts from dolphins with no lung disease diagnosis exhibit broader peaks indicative of the L_α_ phase, fluid nature of these membranes. On the contrary, lung extracts from dolphins with chronic/acute lung disease display a coexistence of sharper peaks, which are indicative of a L_β_ phase, containing closed-packed fatty acid chains. This could be attributed to alterations in acyl chain saturation and/or clustering induced by disease both consistent with the observed increased membrane stiffness (Fig. 4). ***Porras-Gómez et al. (2022b)***

With cryogenic electron microscopy, we observed vesicles (Fig. 5D) in extracts from both diseased (left group) and healthy (right group) lungs displaying high-contrast phospholipid bilayers. The images were analyzed with respect to bilayer thickness and, on average, bilayers from lungs with disease are 1-2 nm thicker than from healthy lung membranes. In line with the presence of L_α_/L_β_ domains ***Heberle et al. (2020)***; ***Cornell et al. (2020)*** we detect two distinct bilayer thicknesses in most vesicles, the L_β_ represented in blue and the L_α_ represented in orange, with the L_β_ domainsbeing roughly 2-3 nm thicker. The main difference between the vesicles from diseased lungs compared to healthy lungs is the size of the vesicle and the fact that larger domains of thicker and stiffer L_β_ (blue) membranes are detected. This is clear by the fact that membranes persist in a straight form (yellow arrows on the left group). This, together with thicker L_β_ domains and a prominent WAXS peak at 15 nm^−1^, is consistent with exacerbated phase separation and membrane stiffening caused by cardiolipin enhancement. Healthy membranes appear more fluid, and a few membrane contacts are apparent between the L_α_ domains of apposed vesicles, which are attributed to the natural process of surfactant protein-driven membrane contacts. ***Parra et al. (2013)*** Indeed, multilamellar polygonal vesicles have been observed by cryo-EM with surfactant protein preferentially located at the membrane bends. ***de la Serna et al. (2013)*** We observe that healthy discrete multilamellar vesicles display an average bilayer separation of *d* ≈ 6.6 nm as well as membrane bends where protein is putatively enriched (Fig. S4). It is also possible that cholesterol facilitates the coexistence of untilted and tilted microdomains that would change the bilayer angle. Indeed, protein-free binary mixtures of DPPC and cholesterol also form polygonal vesicles (Fig. S5). We propose that both cholesterol and surfactant proteins are enriched at the bends of lung membranes.

## Discussion

Supraphysiological amounts of cardiolipins have been detected in lung fluid in terrestrial mammals with pneumonia ***Ray et al. (2010)*** and have been shown to disrupt the structure, mechanical properties, and oxygen permeation of the lipid membrane. ***Steer et al. (2018)***; ***Porras-Gómez et al. (2022b)***; ***Kim et al. (2020)*** In this work, we found analogous alterations in the lipidome, phase behavior, and mechanical and structural properties of lung membranes extracted from clinically affected Navy bottlenose dolphins. This knowledge has the potential to impact future developments of lung disease diagnosis and therapies targeting lung membranes in mammals.

Lipidome analysis shows that both cardiolipins and plasmalogen phosphocholines are more abundant in diseased lungs compared to healthy marine mammal lungs. Plasmalogens, phospholipids associated with inflammatory responses, Bozelli Jr et al. (2021); ***Braverman and Moser (2012)*** are more abundant in lungs with a recent history of disease compared to chronic disease, which positions plasmalogen activity in the early stages of disease. The fact that lung extracts from dolphins with no lung diseases present reduced levels of plasmalogen phosphocholines could be related to the absence of inflammatory triggers. However, further research is required to identify a relation between lung pathology and plasmalogen abundance, and in this paper, we focus on the effects of cardiolipin. While lipidomics clearly suggests that the presence of lung disease significantly alters lipid signatures, our analysis is restricted by the low number of disease-free samples. Lung exudates are collected as part of BAL retrieval, which is not routinely performed in healthy dolphins. Our study mitigates this by employing a multi-prong, robust, and quantitative approach where lipidomics data is backed by several scattering and imaging tools done in multiple replicates per sample, as well as robust statistical analyses.

Alterations in saturation and lipid profiles are expected to affect the phase behavior of membranes. Unsaturated phospholipids tend to form fluid films that are not as efficient at lowering surface tension but present better respreading relative to fully saturated phospholipids. ***Notter (2000)*** Here, we observe that membranes extracted from diseased lungs have less saturation of acyl chains and stronger L_β_ phases, resulting in bigger yet less circular domains. In other words, diseased membranes have less ability to sustain the height mismatch and line tension between the L_β_ and the L_α_ phases. ***García-Sáez et al. (2007)*** Loss of membrane homeostasis has been associated with the enhancement of gel L_β_ phases in senescing cells, ***Ayres (1984)***; ***Thompson et al. (1987)*** during irreversible ischemic cell injury, ***Farber et al. (1981)*** and in erythrocyte cells in cognitive impairment patients. ***Vignini et al. (2019)*** Also, cold-shock injury in cells directly results from aL_α_/L_β_ phase transition in plasma membranes. ***Drobnis et al. (1993)***

Native pulmonary membranes, as the ones used in this study, comprise multiple lipid species, hydrophobic surfactant proteins, and the molecular organization in these domains remains unclear. ***Sear and Cuesta (2003)***; ***Jacobs and Frenkel (2017)*** The L_β_ -L_α_ domains have been reported to align across lipid-stacked multilayers. ***Mills et al. (2008a,b)*** In fact, a reversible monolayer-tomultilayer transition has been reported to occur in pulmonary membranes after further compression. ***Zuo et al. (2008)***; ***Zhang et al. (2011a)***; ***Porras-Gómez et al. (2022a)*** Such transition seems to initiate at the edge of the L_β_ domains where there is an acyl chain mismatch ***Keating et al. (2012)*** and altered elastic and frictional properties. ***Zuo et al. (2008)*** The resultant multilayer structures could provide additional stability, thereby allowing very low surface tensions. ***Zuo et al. (2008)*** This process is reversible, indicating that after squeeze-out, the excluded multilayers remain closely associated with the interfacial monolayer rather than escaping into the hypophase. The excess lipid stored in the multilayer assemblies readily returns into the interface to decrease surface tension. Hence, multilayer structures are required for pulmonary film stability, facilitating low compressibility and low surface tension, and serving as a lipid reservoir for film replenishment. ***Zuo et al. (2008)*** Here, we demonstrate a direct link between lateral membrane heterogeneity and the physiological function of native pulmonary membranes under pathological conditions.

Finally, as expected, lung pathology impacts the mechanical integrity of lung tissue. Pulmonary membranes and tissue from dolphins with no lung disease appear softer because they are composed mostly of discrete membranes separated and cushioned by water layers. In contrast, lung extract membranes from dolphins with chronic/acute lung diseases are dehydrated ***Steer et al. (2018)*** and highly interconnected by hydrophobic cardiolipin-rich stacks, leading to membranes with higher Young modulus ***Porras-Gómez et al. (2022b)*** and stronger adhesion forces. Similar phenomena were observed in porcine pulmonary extracts, where membranes exposed to chemicals known to show acute inhalation toxicity were not able to reach low surface tensions upon compression compared to the control. ***Da Silva et al. (2021)*** The latter correlates with the fact that these same samples exhibit smaller *d*-spacing related to their state of aggregation and dehydration.

Overall, this study advances understanding of how specific lipid alterations fundamentally impact the structure, composition and mechanics of diseased marine mammal lung membranes. Findings from Navy bottlenose dolphins highlight the role of membrane heterogeneity in disease progression and lay the groundwork for improved diagnostic and therapeutic strategies targeting pulmonary membranes in mammals.

## Materials and methods

### Animals

A total of 7 bronchoalveolar lavage (BAL) samples opportunistically collected from U.S. Navy Marine Mammal Program bottlenose dolphins, *Tursiops truncatus*, between February 2017 and April 2022, were analyzed in this study. Samples from Navy animals were collected during their routine care and under the authority codified in U.S. Code, Title 10, Section 7524. Secretary of the Navy Instruction 3900.41H directs that Navy marine mammals be provided the highest quality of care. The U.S. Navy Marine Mammal Program (MMP), Naval Information Warfare Center (NIWC) Pacific, houses and cares for a population of bottlenose dolphins and California sea lions in San Diego Bay (CA, USA). NIWC Pacific is accredited by AAALAC International and adheres to the national standards of the U.S. Public Health Service policy on the Humane Care and Use of Laboratory Animals and the Animal Welfare Act. Five BAL samples were collected opportunistically from animals with naturally occurring respiratory diseases of variable severity. Two BAL samples were also collected opportunistically from the same species without clinical evidence of lung pathology. Based on bacterial and fungal cultures, gross and histopathologic diagnoses, the study included one case of active pneumonia, one case of chronic pulmonary disease of unknown etiology, one case of chronic pulmonary aspergillosis and methicillin-resistant Staphylococcus aureus (MRSA), one case of pulmonary aspergillosis with secondary stenotic airways, one case of suspected *Aspergillus* pneumonia at the time of collection, one case with no prior history of pulmonary disease but with recent ultrasound evidence of alveolar interstitial syndrome, and one case with clinically normal lungs where lung extracts were collected post-mortem. In addition, post-mortem lung tissue samples from 3 dolphins collected during necropsies were included – one sample from a naturally-aborted second-trimester fetus, one sample from a geriatric dolphin with multifocal mild mineralization in the epithelium of the terminal bronchioles and alveolar septa, and one sample from a dolphin with histopathologic evidence of interstitial fibrosis with acute bacterial bronchopneumonia.

### Bronchoalveolar lavage protocol

The raw marine mammal lung exudate was collected via standard bronchoalveolar lavage (BAL) aspiration techniques using a sterilized video endoscope (Karl Storz Slim Gastroscope, outer diameter 5.9 mm, working length 1,100 mm, biopsy channel 2 mm). Briefly, the tip of the bronchoscope was wedged gently in a lung bronchus. A divided, weight-adjusted volume of sterile saline solution (NaCl) was infused, followed by syringe aspiration of the combined fluid and lung exudate. A portion of the recovered fluid and exudate was aliquoted into a large sterile vial and immediately placed into a -80 ^◦^C freezer for pulmonary membrane analysis.

### Lipid extraction

Lipid extraction was performed according to a modified Bligh and Dyer procedure, ***Bligh and Dyer (1959)*** keeping a ratio of 1:1:0.9 (v/v/v) between chloroform/methanol (HPLC grade) and the corresponding aqueous BAL sample. A volume of 200 μL of BAL fluid was placed in a microcentrifuge tube. Volumes of 500 μL of methanol and 250 μL chloroform (chilled to -20 C^◦^) respectively were added. Tubes were mixed using a vortex for 10 sec. The 1-phase solution was left still at 4 C^◦^ for 15-30 minutes. Afterward, 250 μL of ultrapure water and 250 μL of chloroform were added, respectively. The solution, which becomes biphasic at this point, was centrifuged for 10 min at 16000 xg. Finally, the upper polar phase of the solution was discarded, while the bottom organic phase containing the lipid material was transferred to a clean and previously weighed vial. The organic solvent was evaporated in a vacuum desiccator overnight. Weighted dried lipid extracts were stored at -20 C^◦^. Extracts contain all lipid species and hydrophobic surfactant proteins SP-B and SP-C.

### Lipidomics

Dry lipid extracts from BALs were redissolved in a solution of 100 μL of methanol:chloroform (1:2 v/v) and subsequently analyzed by using the Thermo Q-Exactive MS system in the Metabolomics Laboratory of the Roy J. Carver Biotechnology Center, University of Illinois Urbana-Champaign. The mass spectra acquisition protocol was previously reported. ***Zhang et al. (2018)*** The lipid signal responses (peak area counts) were normalized to the corresponding internal standard signals and sample weight. Lipid species were identified by lipid headgroup and fatty acid chain lengths resulting from the generated ions via electrospray ionization (ESI). Since this is based on lipid species affinity to ESI, data are presented as relative abundance, allowing comparison of lipid species between samples. Lipidomics was performed on duplicates and triplicates depending on the sample.

### Lipidomic data analysis

For the lipid analysis, the full data set contained quantitative information from ≈1060 lipid species. Lipids with >80% missing values were filtered out from the analysis (below the detection limit), and the remaining ≈490 lipid species served as the input to subsequent multivariate analyses. Missing values were imputed using the median value of each variable. Data were log-transformed and auto-scaled with mean-centering and divided by the standard deviation of each variable. The top 30 lipids with the highest variable importance in projection (VIP) scores were selected from component 1. Lipid enrichment analysis was performed in LION using 1-way ANOVA F-test P-values or *Log*_2_ fold differences (lung pathology vs.no lung pathology). Distributions of all LION-terms over the ranked list and compared to uniform distributions by using 1-tailed Kolmogorov-Smirnov as previously described. ***Molenaar et al. (2019)*** Enriched LION terms are significant when associated lipids are higher ranked than expected by chance. Z-scores are averages ± SEM and analyzed with Kruskal-Wallis test (*p<0.05). Lipid species-level analysis of cardiolipins was performed by using *Log*_2_ fold differences and FDR *q*-values (chronic/acute lung disease vs. no lung disease).

### Atomic Force Microscopy

BAL extracts were dissolved in chloroform and afterward spin-coated onto 1 cm^2^ thermal oxide SiO_2_ substrates (University Wafer Inc., MA, USA) at 4000 RPM for 30 s using a spin coater (VTC-100A, MTI Corporation, California, USA). The samples were vacuum-dried for at least 2 hours and scanned in air afterward using a high-speed Cypher AFM (Asylum Research-Oxford Instruments, California, USA). Contact mode AFM probes (ContE-G, *K*= 0.2 N/m) with platinum overall coating were used (NanoAndMore U.S.A. Corporation, California, USA). Lung tissue from dolphins kept at -80 °C was snap-frozen after submerging directly in isopentane in a liquid nitrogen bath for about 15 s. Afterward, the frozen tissue was embedded in an optimal cutting temperature (OCT) compound. The tissue was then cryo-cut into 5 μm sections onto poly-L-lysine-coated ***Wenderott et al. (2020)*** thermal oxide SiO_2_ substrates (2.5 cm^2^). To ensure the AFM tip scanned the tissue and not the OCT, only one slide was deposited in the center of a substrate, making it visually easier to locate the tissue sample. Phosphate-buffered saline (PBS) was added on top of the sample and on the AFM tip before bringing them into contact. We used gold-coated (on both sides) silicon nitride cantilevers (HYDRA6R-200NGG, AppNano, Mountain View, CA) with a nominal spring constant of 0.035 N/m. Gold coating on both sides decreases adhesion and improves tip-tissue interaction. For both BAL extracts and tissue samples, the cantilever spring constant and the resonance frequency were calibrated using the thermal spectrum before each experiment. The amplitude sensitivity was calibrated in air and in fluid (for tissue). The force applied to the samples was fixed to the lowest possible value, allowing reproducible mapping (1 V corresponding to a force range of 1-2 nN). The scan rate was set to 100 Hz on an area of 5 x 5 μm. The total *Z*-piezo (vertical) displacement was set to up to 2 μm. Force maps comprised 128 points x 128 lines, resulting in 16,384 data points per map. Force maps were processed using the accompanying Asylum Research software v.16. All reported Young modulus *E* values resulted from the peak of a Gaussian fit of *E* distribution obtained from the respective force maps. AFM force mapping was performed in a high-speed Cypher AFM (Asylum Research Oxford Instruments, California, USA).

### Circularity analysis

To study the effect of phase separation in pulmonary extracts, the domains were characterized by their area and circularity. Open-source ImageJ software (National Institute of Health, USA) was used to obtain quantitative circularity and domain area. Circularity was calculated as 4π×area/(perimeter)^2^ for every measured domain. Statistical analyses were performed using JMP software.

### Confocal microscopy

For confocal microscopy, lipid extracts were fluorescently labeled by incorporating Rhodamine lipids (16:0 and 18:1 Liss Rhod PE) in a total concentration of 0.1 mol% and imaged in an LSM 800 confocal microscope (Carl Zeiss 407 Microimaging, Jena, Germany). For TEM imaging, BAL lipid extracts were rehydrated at 2 mg/mL and incubated at 55 ^◦^C for 24 h. After negative staining, ***Baxa (2018)*** 5 μL were deposited on Holey Carbon supported Copper grids, size 200 mesh, size 100 nm (Sigma Aldrich) and imaged in a JEM-1400 transmission electron microscope (JEOL USA Inc).

### X-ray scattering

X-ray analysis detailed protocol can be found in previous work. ***Rueben et al. (2022)*** Briefly, dried lipid extracts were rehydrated into quartz capillaries to a final concentration of 45 mg/mL. Samples were analyzed using Synchrotron SAXS and WAXS at beamline 12-ID-B of the Advanced Photon Source at Argonne National Laboratory. For peak area comparisons, peaks were analyzed using the Peak Analyzer tool included in OriginPro 2019. Scattering data from water in the capillary was subtracted from all WAXS sample data to enhance WAXS peak intensity.

### Transmission electron microscopy

BAL lipid extracts were rehydrated at 2 mg/mL and incubated at 55 ^◦^C for 24 h. After negative staining, ***Baxa (2018)*** 5 μL were deposited on Holey Carbon supported Copper grids, size 200 mesh, size 100 nm (Sigma Aldrich) and imaged in a JEM-1400 transmission electron microscope (JEOL USA Inc). Tissue samples were fixed overnight at 4°C in 2.5% (w/v) paraformaldehyde and 2% (w/v) glutaraldehyde dissolved in 0.2 M phosphate buffer, pH 7.2. They were then incubated in 2% (w/v) osmium tetroxide (EMS, 19170) for 1 hour at ambient temperature. A volume of 5% (w/v) potassium ferrocyanide (EMS, 25154-5) was then added to the samples and incubated for an additional 15 minutes. After extensive washes in water, samples were *en bloc* stained with saturated uranyl acetate (JT Baker Chemicals, 4192) in water and dehydrated in ascending concentrations of ethanol until the tissues were in 100% ethanol. Ethanol was replaced with pure acetonitrile, and samples were incubated for 10 minutes. Samples were then infiltrated with ascending concentrations of LX112 resin (LADD, 21210) in acetonitrile mixtures. Finally, samples were embedded in 100% LX112 resin and polymerized at 60°C overnight. Embedded samples were then cut into 70 nm sections using a diamond knife mounted on a Leica UC6 Ultramicrotome. Sections were collected on 200 mesh holey carbon copper grids (SPI Supplies, 3620C-MB) and imaged using a JEOL JEM-1400 transmission electron microscope.

### Cryogenic electron microscopy

BAL organic extracts rehydrated in water and whole BAL aqueous samples for cryogenic BAL organic extracts rehydrated in water and whole BAL aqueous samples for cryogenic transmission electron microscopy were prepared on a QUANTIFOIL R 2/1 copper mesh grid (Electron Microscopy Sciences) using a Vitrobot Mark IV (Thermo Fisher Scientific). A volume of 3.5 μL of sample was applied to the glow-discharged surface of the grid at a chamber temperature of 6 ^◦^C with a relative humidity level of 100%, and then vitrified in liquid ethane. Before vitrification, samples were blotted for 6 seconds with a blot force of 6. All Cryo-EM grids were imaged using a 200 keV Glacios (Thermo Fisher Scientific) TEM equipped with a Falcon 4 direct electron detector. Images were collected using EPU automated acquisition software (Thermo Fisher). Additional qualitative analysis was conducted using FIJI software ***Schindelin et al. (2012)*** to perform thickness measurements on a single bilayer and to evaluate the distance between adjacent lamellae. The thickness measurements were taken by evaluating the distance between the outer boundaries of the high-contrast bilayer regions present in the images. The average interlamellar spacing values were obtained using the inverted FFT method, usually implemented to calculate d-spacing values from the crystalline structure of individual nanoparticles. By implementing this analysis on cryo-EM images containing clustered bilayers, the electron intensity profiles were generated. Subsequently, peak-to-peak measurements were extracted from all electron profiles to obtain the average interlamellar spacings.

## Acknowledgments

This work was funded by the Office of Naval Research (ONR) grant no. N00014-21-1-2029 and used a confocal microscope acquired with support from an ONR DURIP (Defense University Research Instrumentation Program) Grant Number N00014-18-1-2087. This research used resources of the Advanced Photon Source, a U.S. Department of Energy (DOE) Office of Science user facility operated for the DOE Office of Science by Argonne National Laboratory under Contract No. DE-AC02-06CH11357. This research was carried out in part in the Materials Research Laboratory, Central Research Facilities, University of Illinois. The lipidomics work was partially supported by the Cancer Center at the University of Illinois Urbana-Champaign. LC-MS-based lipidomic analysis was performed at the Roy J. Carver Biotechnology Center, University of Illinois Urbana-Champaign. Tissue slicing for AFM was carried out by Kingsley Boateng in the Core Facilities at the Carl R. Woese Institute for Genomic Biology, University of Illinois Urbana-Champaign.

